# Molecular dynamics analysis of Superoxide Dismutase 1 mutations suggests decoupling between mechanisms underlying ALS onset and progression

**DOI:** 10.1101/2022.12.05.519128

**Authors:** Munishikha Kalia, Mattia Miotto, Deborah Ness, Sarah Opie-Martin, Thomas P Spargo, Lorenzo Di Rienzo, Tommaso Biagini, Francesco Petrizzelli, Ahmad Al-Khleifat, Renata Kabiljo, Simon Topp, Keith Mayl, Isabella Fogh, Puja R Mehta, Kelly L Williams, Jennifer Jockel-Balsarotti, Taha Bali, Wade Self, Lyndal Henden, Garth A Nicholson, Nicola Ticozzi, Diane McKenna-Yasek, Lu Tang, Pamela Shaw, Adriano Chio, Albert Ludolph, Jochen H Weishaupt, John E Landers, Jonathan D Glass, Jesus S Mora, Wim Robberecht, Philip Van Damme, Russell McLaughlin, Orla Hardiman, Leonard H van den Berg, Jan H Veldink, Phillippe Corcia, Zorica Stevic, Nailah Siddique, Antonia Ratti, Vincenzo Silani, Ian P Blair, Dong-sheng Fan, Florence Esselin, Elisa de la Cruz, William Camu, A Nazli Basak, Teepu Siddique, Timothy Miller, Robert H Brown, Peter M Andersen, Project MinE ALS Sequencing Consortium, Christopher E Shaw, Tommaso Mazza, Giancarlo Ruocco, Edoardo Milanetti, Richard JB Dobson, Ammar Al-Chalabi, Alfredo Iacoangeli

## Abstract

Mutations in the superoxide dismutase 1 (*SOD1*) gene are the second most common known cause of ALS. *SOD1* variants express high phenotypic variability and over 200 have been reported in people with ALS. Investigating how different *SOD1* variants affect the protein dynamics might help in understanding their pathogenic mechanism and explaining their heterogeneous clinical presentation. It was previously proposed that variants can be broadly classified in two groups, ‘wild-type like’ (WTL) and ‘metal binding region’ (MBR) variants, based on their structural location and biophysical properties. MBR variants are associated with a loss of SOD1 enzymatic activity. In this study we used molecular dynamics and large clinical datasets to characterise the differences in the structural and dynamic behaviour of WTL and MBR variants with respect to the wild-type SOD1, and how such differences influence the ALS clinical phenotype. Our study identified marked structural differences, some of which are observed in both variant groups, while others are group specific. Moreover, applying graph theory to a network representation of the proteins, we identified differences in the intramolecular contacts of the two classes of variants. Finally, collecting clinical data of approximately 500 *SOD1* ALS patients carrying variants from both classes, we showed that the survival time of patients carrying an MBR variant is generally longer (~6 years median difference, p < 0.001) with respect to patients with a WTL variant. In conclusion, our study highlights key differences in the dynamic behaviour of the WTL and MBR SOD1 variants, and wild-type SOD1 at an atomic and molecular level. We identified interesting structural features that could be further investigated to explain the associated phenotypic variability. Our results support the hypothesis of a decoupling between mechanisms of onset and progression of *SOD1* ALS, and an involvement of loss-of-function of SOD1 with the disease progression.

## INTRODUCTION

Amyotrophic lateral sclerosis (ALS) is a fatal neurodegenerative disorder, characterized by progressive muscle weakness and paralysis, leading to death from neuromuscular respiratory failure typically within 5 years of symptom onset [1, 2]. Mutations in the *superoxide dismutase type 1 (SOD1)* gene are the second most common known cause of ALS and have been found in both familial ALS and sporadic ALS [3]. SOD1 is an antioxidant metalloenzyme, which protects cells from oxidative damage. It catalyses the conversion of the superoxide O_2_^-^ free radical present in the cytoplasm to H_2_O_2_ and molecular oxygen. SOD1 is a 32 kDa protein which exists as a homodimer and its structure is characterized by the presence of a β-barrel immunoglobin fold, two long loops: metal binding loop (MBL) and electrostatic loop (ESL) [4]. The fully formed SOD1 homodimer is highly stable and has a melting point of 92 °C [5], due to copper and zinc binding, an intrasubunit disulphide bond and dimerization [6].

At present, more than 220 *SOD1* mutations have been reported in people with ALS [7, 8]. Although there is evidence that *SOD1* mutations cause ALS, the exact underlying mechanism of disease onset and progression remains poorly understood [9]. It has been proposed that the pathogenic mutations hinder post-translational maturation, which decreases structural stability, thereby instigating the misfolding of SOD1 protein [10–12]. The ultimate outcome of SOD1 misfolding is the formation of hallmark amyloid aggregates of SOD1 in the affected tissues [13–15]. A noteworthy feature of the misfolded SOD1 is its prion-like behaviour of self-proliferation. It was proposed that a misfolded SOD1 can transverse between cells and cause misfolding of other SOD1 molecules via the protein aggregates released from dying cells [16–18].

Moreover, *SOD1* variants can express high phenotypic variability. For example, A4V (A5V in the HGVS V2.0 nomenclature – traditional nomenclature will be used throughout this manuscript) is a variant which is responsible for 48% of *SOD1* ALS in US familial ALS. It is associated with variable site of onset, rapid disease progression and a mean survival time of 1.1 years from clinical presentation [19–21]. In contrast, H46R is commonly observed in the Japanese, Pakistan and Norwegian populations and is invariably associated with a stereotypic phenotype with slowly ascending paresis beginning in the legs and long survival (approx. 12 years from diagnosis) [22–25]. Investigating the differences between *SOD1* mutations in terms of their structural effect on the protein might help in understanding the mechanism underlying such a heterogeneous clinical presentation.

Disease-associated *SOD1* mutations are found in all domains of the protein and over the years attempts have been made to classify the impact of each variant based on location and subsequent effect on protein structure [26]. Based on their structural location and biophysical characteristics such as metal binding affinity and influence on SOD1 *in vitro* activity, it has been proposed that *SOD1* variants can be classed in two broad groups [26–28]. The first group consists of the ‘wild-type like’ (WTL) variants such as: A4V, L38V, G37R, G41S, G72S, D76Y, D90A and G93A. These variants bind metal ions tightly and the catalytic activity is not affected *in vitro*, therefore their biophysical behaviour is expected to be similar to that of the wild-type SOD1 (wt-SOD1) [27, 29, 30]. The second group contains ‘metal binding region’ (MBR) variants, such as: H46R, H48Q, G85R, D124V, D125H, G127X and S134N. These variants are localized in the metal binding sites, metal binding loop and electrostatic loop regions, and cause reduced metal binding and diminished *in vitro* catalytic activity [4, 31–33].

The aim of this study was to characterize the differences in the structural and dynamic behaviour of the WTL and MBR disease-causing SOD1 variants with respect to the wt-SOD1. For this we performed all atomistic molecular dynamics (MD) simulations of the wt-SOD1, 6 MBR and 7 WTL SOD1 variant monomers. We focussed on the metal-free SOD1 monomer, as the conformational changes of the metal binding loop and electrostatic loop are unravelled by the lack of metal stabilisation [33]. In order to quantify and characterise the motion of the whole protein and its loops, we studied root mean square deviation (RMSD), root mean square fluctuation (RMSF), radius of gyration, number of hydrogen bonds and performed principal component analysis (PCA). Moreover, two additional analyses were carried out that focus on the study of individual residues. In the first we studied the covariance of the residue motions and in the second, we used a graph-theory based approach to study the complexity of the intermolecular connections that were captured in a simple residue-specific descriptor. Our results suggested that the structural and dynamic behaviour differences between WTL and MBR variants could be linked to the ALS clinical phenotype. To explore this hypothesis, we collected a large clinical dataset of *SOD1* ALS patients [34] and performed survival and age of onset analyses in the patients carrying WTL and MBR variants.

## RESULTS

To characterize the impact of one mutation at a time, we focussed only on SOD1 structures with one single amino acid variant. We could identify 13 WTL and MBR mutations in the Protein Data Bank (PDB) for which high quality structures were available. We divided them in the two categories based on whether they were previously reported to belong to one or whether they were in proximity of another mutation of known category. For example, we classified T2D as WTL as it is in proximity of A4V which was reported to exhibited normal SOD activity *in vitro* [27]. The WTL group included T2D, A4V, G37R, L38V, T54R, G93R, I113T and the MBR variants were H46R, C57S, H80R, G85R, D124V and D125H. The structural location of the variants is shown in the Figure 1. The detailed structural information, clinical interpretation, and resolution of the PDB structures used in the study are shown in Table 1.

**Figure 1.**
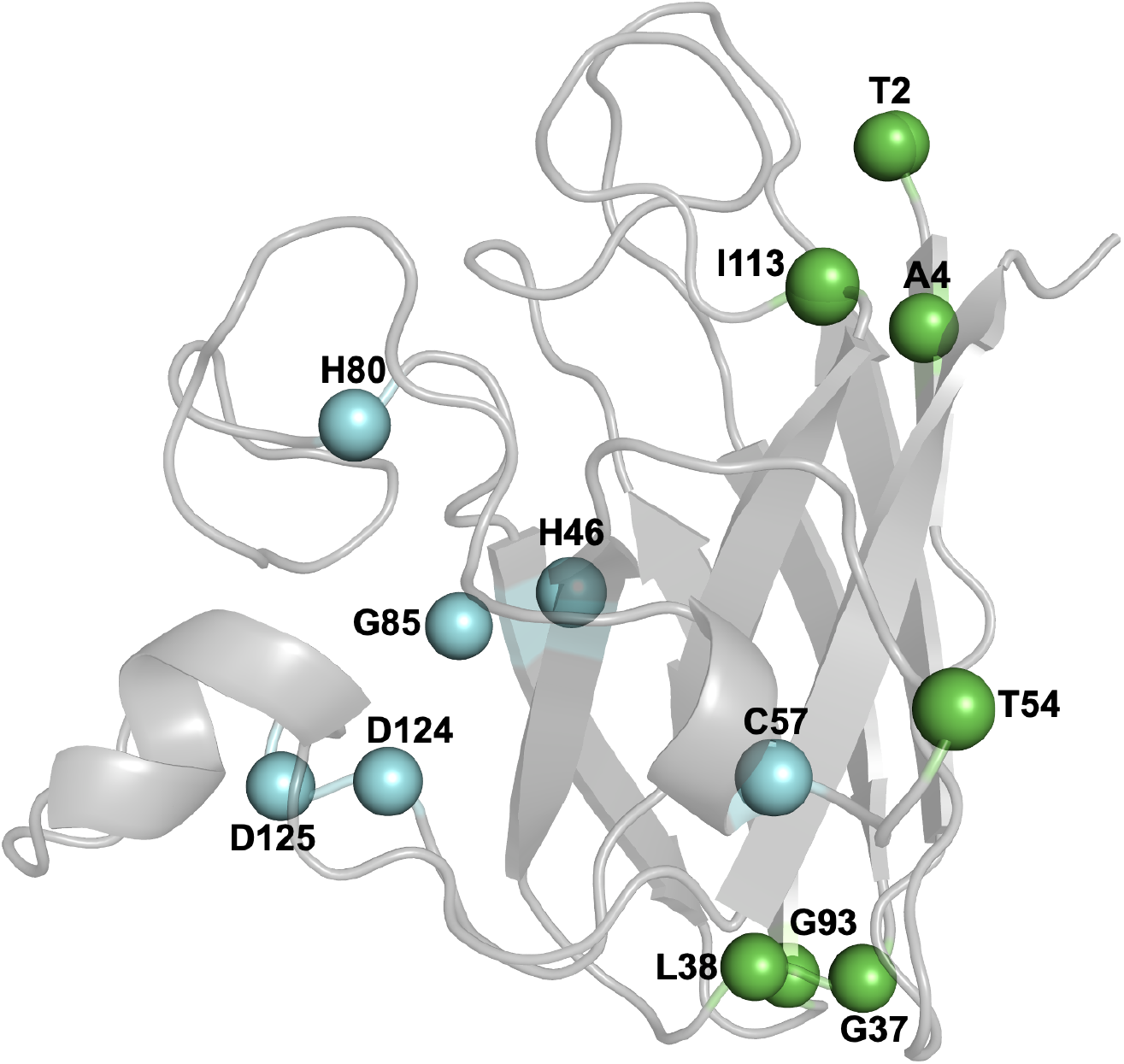
SOD1 monomer structure coloured in grey, the residues which form WTL variants are coloured in blue and the residues which form the MBR variants are coloured in green.

**Table 1:**
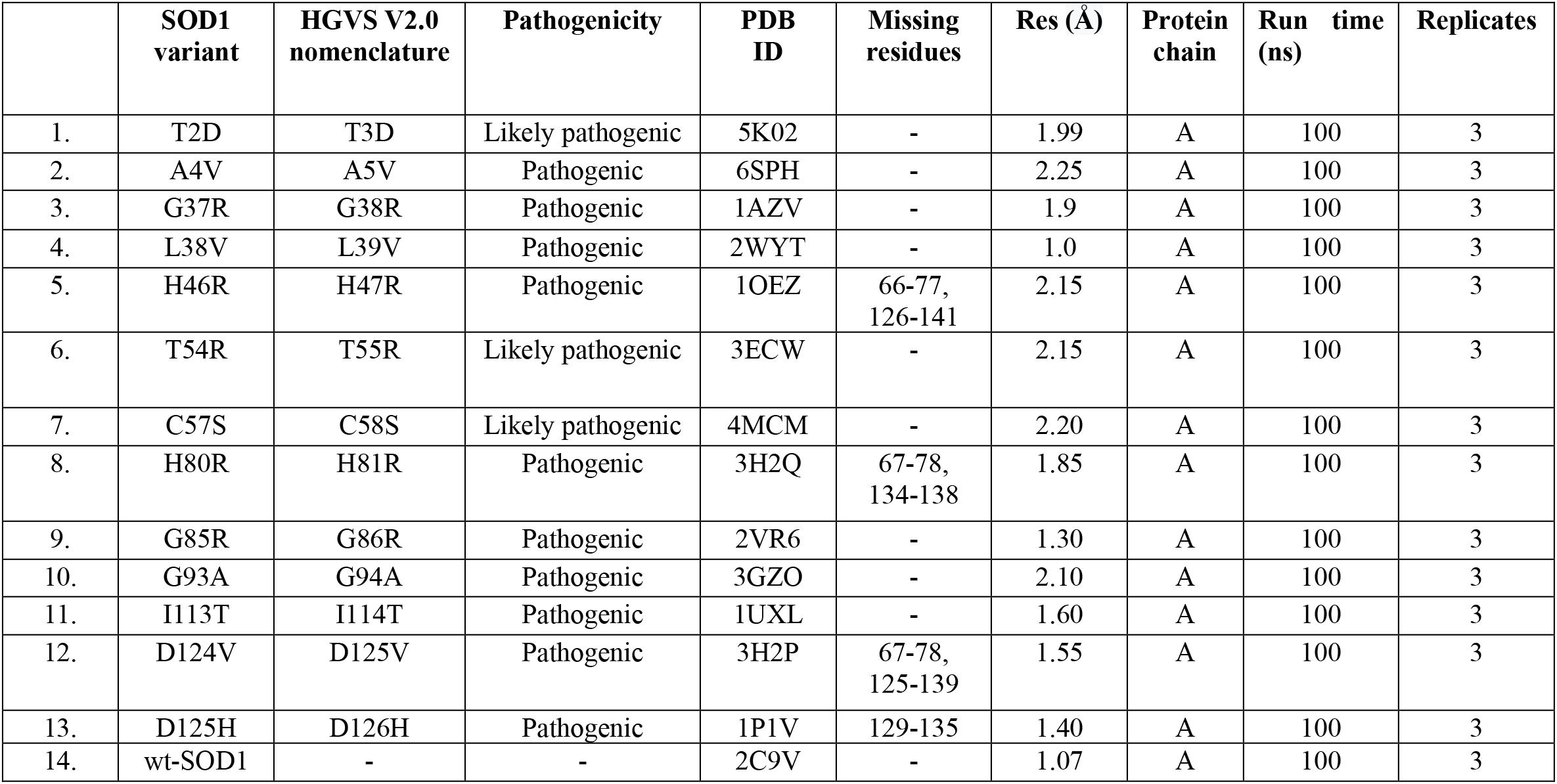
List of PDB structures of the wt-SOD1 and its variants used in the study.

|  | SOD1 variant | HGVS V2.0 nomenclature | Pathogenicity | PDB ID | Missing residues | Res (Å) | Protein chain | Run time (ns) | Replicates |
| --- | --- | --- | --- | --- | --- | --- | --- | --- | --- |
| 1. | T2D | T3D | Likely pathogenic | 5K02 | - | 1.99 | A | 100 | 3 |
| 2. | A4V | A5V | Pathogenic | 6SPH | - | 2.25 | A | 100 | 3 |
| 3. | G37R | G38R | Pathogenic | 1AZV | - | 1.9 | A | 100 | 3 |
| 4. | L38V | L39V | Pathogenic | 2WYT | - | 1.0 | A | 100 | 3 |
| 5. | H46R | H47R | Pathogenic | 1OEZ | 66-77, 126-141 | 2.15 | A | 100 | 3 |
| 6. | T54R | T55R | Likely pathogenic | 3ECW | - | 2.15 | A | 100 | 3 |
| 7. | C57S | C58S | Likely pathogenic | 4MCM | - | 2.20 | A | 100 | 3 |
| 8. | H80R | H81R | Pathogenic | 3H2Q | 67-78, 134-138 | 1.85 | A | 100 | 3 |
| 9. | G85R | G86R | Pathogenic | 2VR6 | - | 1.30 | A | 100 | 3 |
| 10. | G93A | G94A | Pathogenic | 3GZO | - | 2.10 | A | 100 | 3 |
| 11. | I113T | I114T | Pathogenic | 1UXL | - | 1.60 | A | 100 | 3 |
| 12. | D124V | D125V | Pathogenic | 3H2P | 67-78, 125-139 | 1.55 | A | 100 | 3 |
| 13. | D125H | D126H | Pathogenic | 1P1V | 129-135 | 1.40 | A | 100 | 3 |
| 14. | wt-SOD1 | - | - | 2C9V | - | 1.07 | A | 100 | 3 |

We have used the monomeric apo SOD1 instead of the fully formed SOD1 homodimer, as it is the least stable form of the protein and an ideal model to study the impact of the pathogenic variants [32, 33]. We performed all atomistic MD simulations of the wt-SOD1, 6 MBR and 7WTL variants in TIP3P water for 100 ns each and each simulation was performed in triplicates.

### Flexibility of the metal binding and electrostatic loops

The root mean square fluctuation is a measure of individual protein residue flexibility. A high RMSF indicates higher flexibility and a low RMSF indicates lower flexibility. We analysed the RMSF of Cα atoms of the wt-SOD1 and 13 SOD1 variants. Our analysis highlighted marked differences in the flexibility of the metal binding loop and electrostatic loop regions (Figure 2a, b), while the remaining part of the protein was relatively less flexible (Sup Fig 1). For the metal binding loop region (Figure 2a), the highest flexibility was observed in the H46R variant. Next, we performed the Wilcoxon Rank-Sum Test for statistical comparison between the RMSF of each variant and the wt-SOD1. We noted that the RMSF difference between a variant and the wt-SOD1 was significant for A4V, H46R and D125H. For the remaining variants the RMSF difference between the variants and the wt-SOD1 were non-significant.

**Figure 2.**
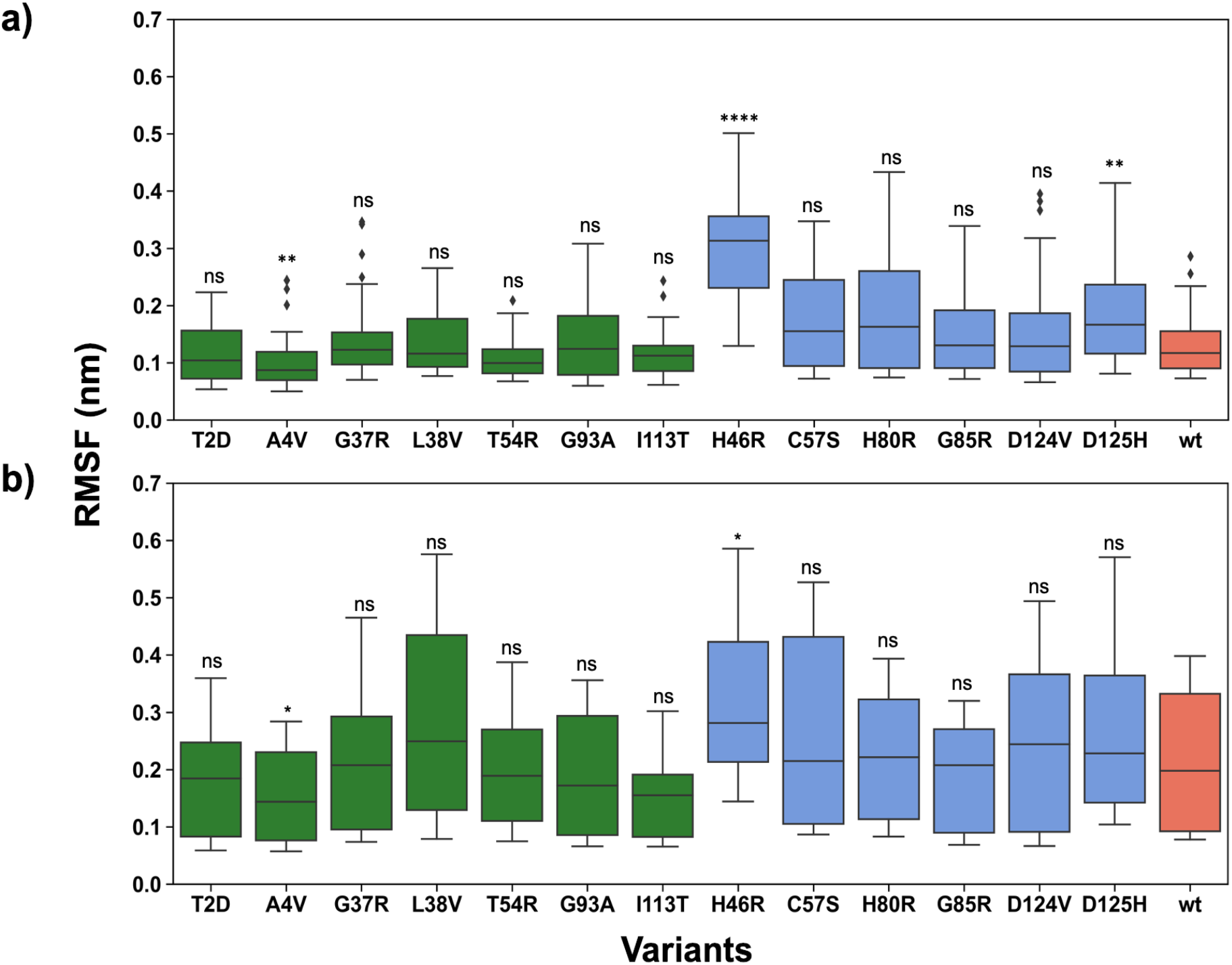
Boxplots depicting the RMSF distribution of the WTL (green), MBR (blue) variants and wt-SOD1 (coral) in the a) MBL b) ESL regions. RMSF is calculated for the Cα atoms. The whisker bars represent the range of minimum and maximum RMSD, the median is represented by a line subdividing the box. Wilcoxon rank-sum test, p-value annotation legend: ns: p <= 1.00e+00, *: 1.00e-02 < p <= 5.00e-02, **: 1.00e-03 < p <= 1.00e-02, ***: 1.00e-04 < p <= 1.00e-03, ****: p <= 1.00e-04.

Focussing on the electrostatic loop, the MBR variants such as C57S, D125H, H46R H80R (Figure 2b), generally showed higher RMSF with respect to WTL variants, illustrating their increased flexibility. In contrast, WTL variants, such as A4V, G37R, I113T and G93A, showed FMSF values lower or very similar to that of wt-SOD1 (Figure 2b). The only exception was L38V which had very high RMSF values in the electrostatic loop region. The statistical analysis showed that the RMSF difference between A4V, H46R and wt-SOD1 were significant, while for the remaining variants the differences were not significant. A4V causes one of the most aggressive forms of *SOD1* ALS, whilst H46R results in very slow progression. It is interesting to note that their RMSFs corresponded to the lowest and highest values among the investigated variants, suggesting that such dynamics features could be explored to explain the clinical representation of variants.

Finally, we tested the difference between the RMSF distributions of the entire protein for all the variants in each class, i.e. WTL and MBR, and the wt-SOD1 (Sup Fig 2). The RMSF of the WTL variants was not different from the wt-SOD1 (p = 0.54) while the RMSF of the MBR variants was higher than both the wt-SOD1 (p = 5.2×10^-07) and the WTL variants (p = 6.4×10^-08).

### Comparison of the conformational changes in the WTL and MBR variants

The root mean square deviation is a quantitative measure of the similarity between two superimposed sets of atomic coordinates and is widely used in the MD analysis to highlight the conformational changes observed in the protein structures over the length of simulations. We observed that there are clear differences in the Cα RMSD distribution of the WTL and MBR variants (Figure 3 and Table 2). The RMSD distributions of the WTL variants were similar to that of the wt-SOD1. The mean and median RMSDs observed in all WTL variants were <= than the mean and median RMSD of the wt-SOD1 (both 0.22 nm), while all mean and median RMSDs observed for all MBR variants but G85R, for which the mean was 0.20 nm and median 0.21nm, were higher than the wt-SOD1 and WTL variants. The same trend was observed when only the RMSDs between 20ns and 100ns were considered to account for the equilibration phase (Sup Fig 3).

**Figure 3.**
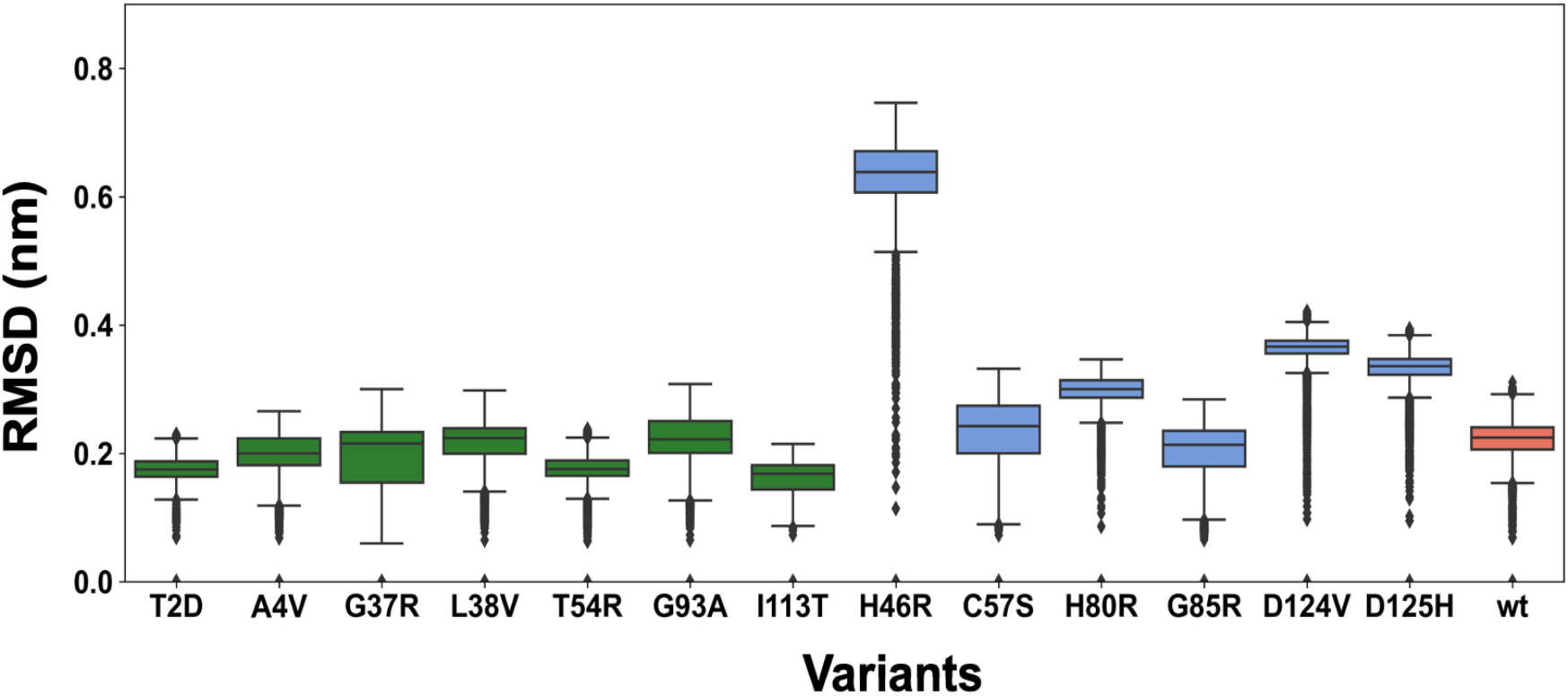
Box plots depicting RMSD analysis of the MD simulations performed on the wt-SOD1 (coral), WTL (green) and MBR (blue) variants. The RMSD is calculated for the Cα atoms of the wt-SOD1 and the variants. The whisker bars represent the range of minimum and maximum RMSD, the median is represented by a line subdividing the box.

**Table 2.** The mean and median of the RMSD calculated for the WTL, MBR variants and wt-SOD1.

|  | **SOD1 variant** | **Mean (nm)** | **Median (nm)** |
| --- | --- | --- | --- |
| 1. | T2D | 0.18 | 0.17 |
| 2. | A4V | 0.20 | 0.20 |
| 3. | G37R | 0.20 | 0.22 |
| 4. | L38V | 0.22 | 0.22 |
| 5. | T54R | 0.17 | 0.18 |
| 6. | G93A | 0.22 | 0.22 |
| 7. | I113T | 0.16 | 0.17 |
| 8. | H46R | 0.63 | 0.64 |
| 9. | C57S | 0.23 | 0.24 |
| 10. | H80R | 0.30 | 0.30 |
| 11. | G85R | 0.20 | 0.21 |
| 12. | D124V | 0.36 | 0.37 |
| 13. | D125H | 0.33 | 0.34 |
| 14. | wt | 0.22 | 0.22 |

Subsequently, we investigated the conformational changes adopted by the variants and the wt-SOD1 during the simulations. We observed that the wt-SOD1 adopted one stable conformation during the simulations (Figure 4). Among the WTL variants, L38V and T54R also assumed a single conformation. A4V, G93A and I113T acquired two major conformations, whilst T2D and G37R adopted three major conformations. The MBR variants D124V and D125H presented one major conformation for the entire length of the simulation, H46R and H80R adopted two major conformations and, interestingly, C57S and G85R variants showed four major conformations during the trajectory of the simulations (Figure 4).

**Figure 4.**
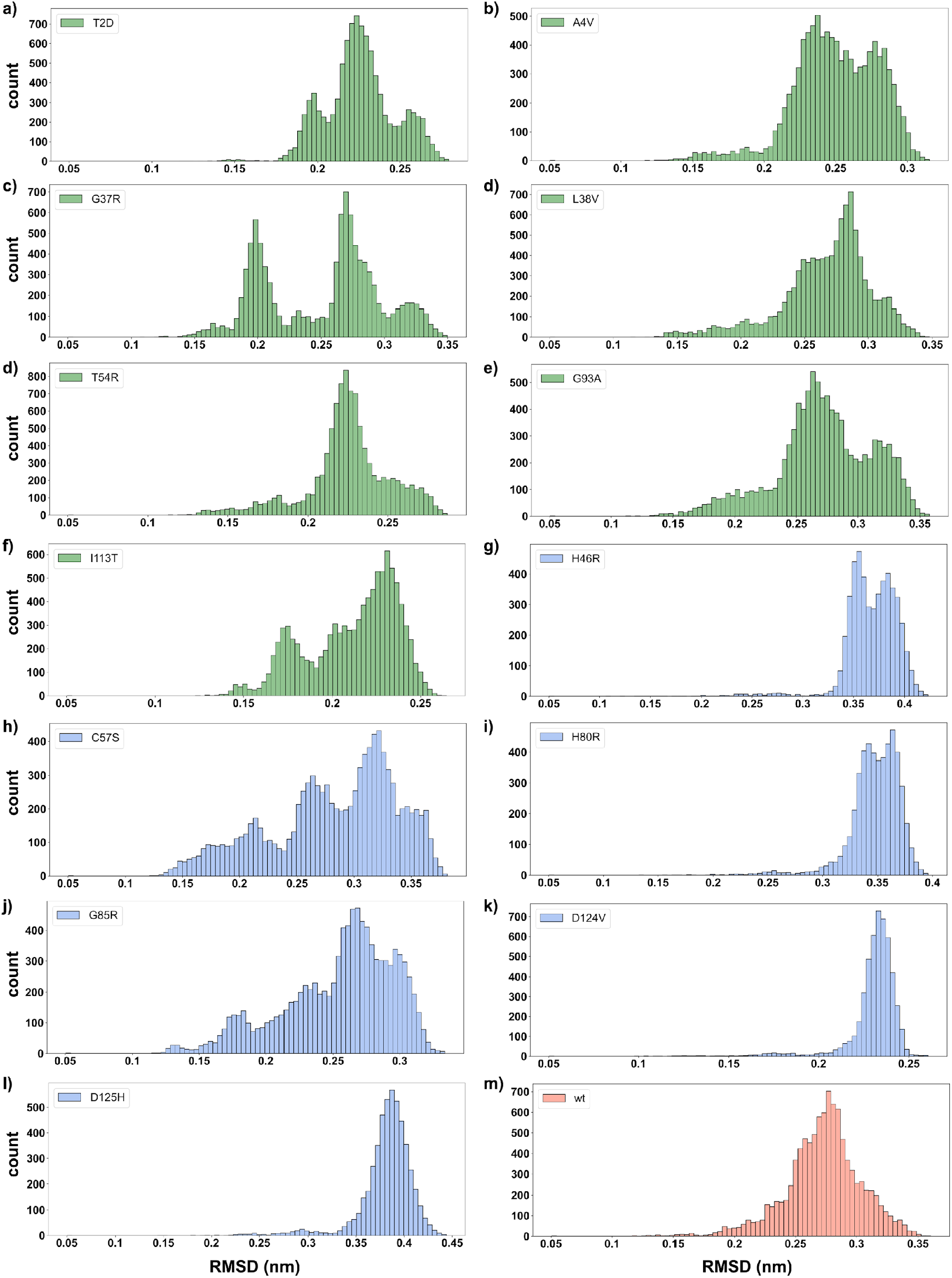
Histograms showing the conformational changes observed in the WTL (green), MBR (blue) variants and wt-SOD1 (coral). a) T2D b) A4V c) G37R d) L38V e) G93A f) I113T g) H46R h) C57S i) H80R j) G85R k) D124V l) D125H m) wt-SOD1.

### Comparison of the compactness of the WTL and MBR variants

The radius of gyration (Rg) is a measure of the compactness of a protein and is widely utilised to compare the protein’s dynamic behaviour. A high radius of gyration is an indication that the protein structure is more extended, whilst a smaller radius of gyration is indicative of a more compact protein structure. We calculated the radius of gyration of the WTL and MBR variants, and wt-SOD1 over the entire length of the simulations. Our analysis showed that the protein structure remains closely packed in the wt-SOD1 and WTL variants (Figure 5), while MBR variants adopt a more extended conformation (Figure 5). The largest difference with respect to the wt-SOD1 was observed in the H46R and H80R variants.

**Figure 5.**
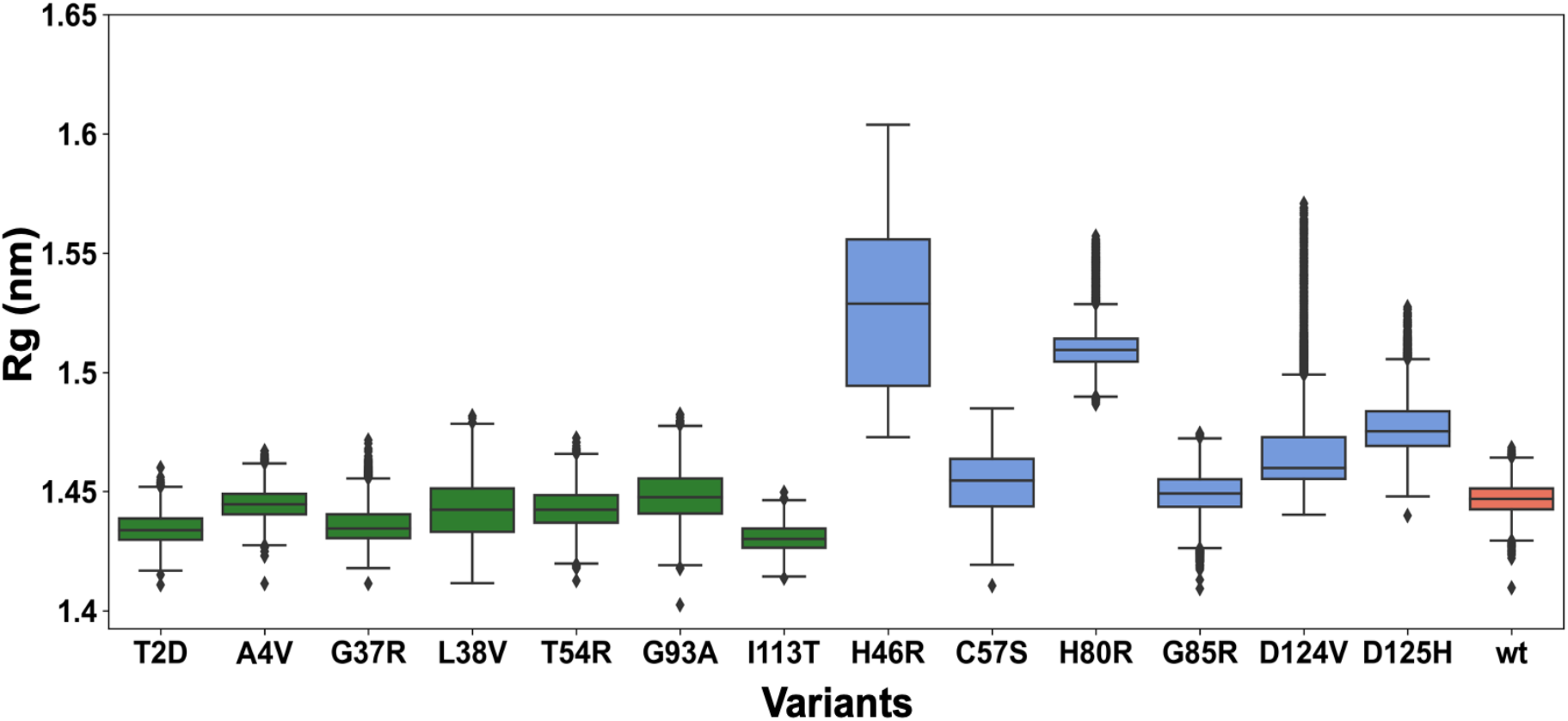
Box plots depicting radius of gyration (Rg) comparison of the wt-SOD1 (coral), WTL (green) and MBR (blue) variants simulations. The whisker bars represent the range of minimum and maximum Rg, the median is represented by a line subdividing the box.

### Principal Component Analysis

Principal Component Analysis (PCA) is a multivariate statistical technique that is frequently used to reduce the number of dimensions that describe the dominant protein domain motion [35–37]. PCA was performed on the WTL, MBR variants and wt-SOD1. The first three eigenvectors explained > 40 % of all the domain motion (Table 3) and such motion was driven by the electrostatic loop for the wt-SOD1 and all variants but T2D and I113T (porcupine plots in Figure 6 a,b). In the MBR SOD1 variants, considerable domain motion was also observed in the metal binding loop in addition to the electrostatic loop. Moreover, the magnitude of the domain motion in the metal binding loop and electrostatic loop regions was substantially larger for the MBR variants than the WTL variants. It was interesting to note that the dynamic motion of metal binding loop and electrostatic loop regions in L38V was more similar to the MBR variants than the WTL variants. The motion variance explained by PC1-4 for each variant and the wt-SOD1 is shown in Table 3.

**Table 3.** Percentage of variance represented by PC1, PC2, PC3, PC4.

|  | **SOD1 variant** | **PC1 (%)** | **PC2 (%)** | **PC3 (%)** | **PC4 (%)** |
| --- | --- | --- | --- | --- | --- |
| 1. | A4V | 28.93 | 19.89 | 9.31 | 6.25 |
| 2. | C57S | 35.31 | 24.03 | 5.63 | 3.44 |
| 3. | D124V | 30.17 | 20.63 | 5.46 | 4.97 |
| 4. | D125H | 30.88 | 15.22 | 9.25 | 5.71 |
| 5. | G37R | 51.56 | 12.36 | 5.74 | 3.24 |
| 6. | G85R | 37.39 | 10.61 | 5.84 | 4.49 |
| 7. | G93A | 22.21 | 13.09 | 11.35 | 7.04 |
| 8. | H46R | 36.99 | 25.66 | 6.52 | 4.24 |
| 9. | H80R | 29.2 | 13.27 | 9.06 | 5.21 |
| 10. | I113T | 33.31 | 20.78 | 8.04 | 5.14 |
| 11. | L38V | 40.32 | 18.38 | 7.23 | 5.29 |
| 12. | T2D | 32.74 | 9.58 | 6.5 | 5.22 |
| 13. | T54R | 27.44 | 13.15 | 9.17 | 5.71 |
| 14. | wt-SOD1 | 27.45 | 21.57 | 10.59 | 4.05 |

**Figure 6.**
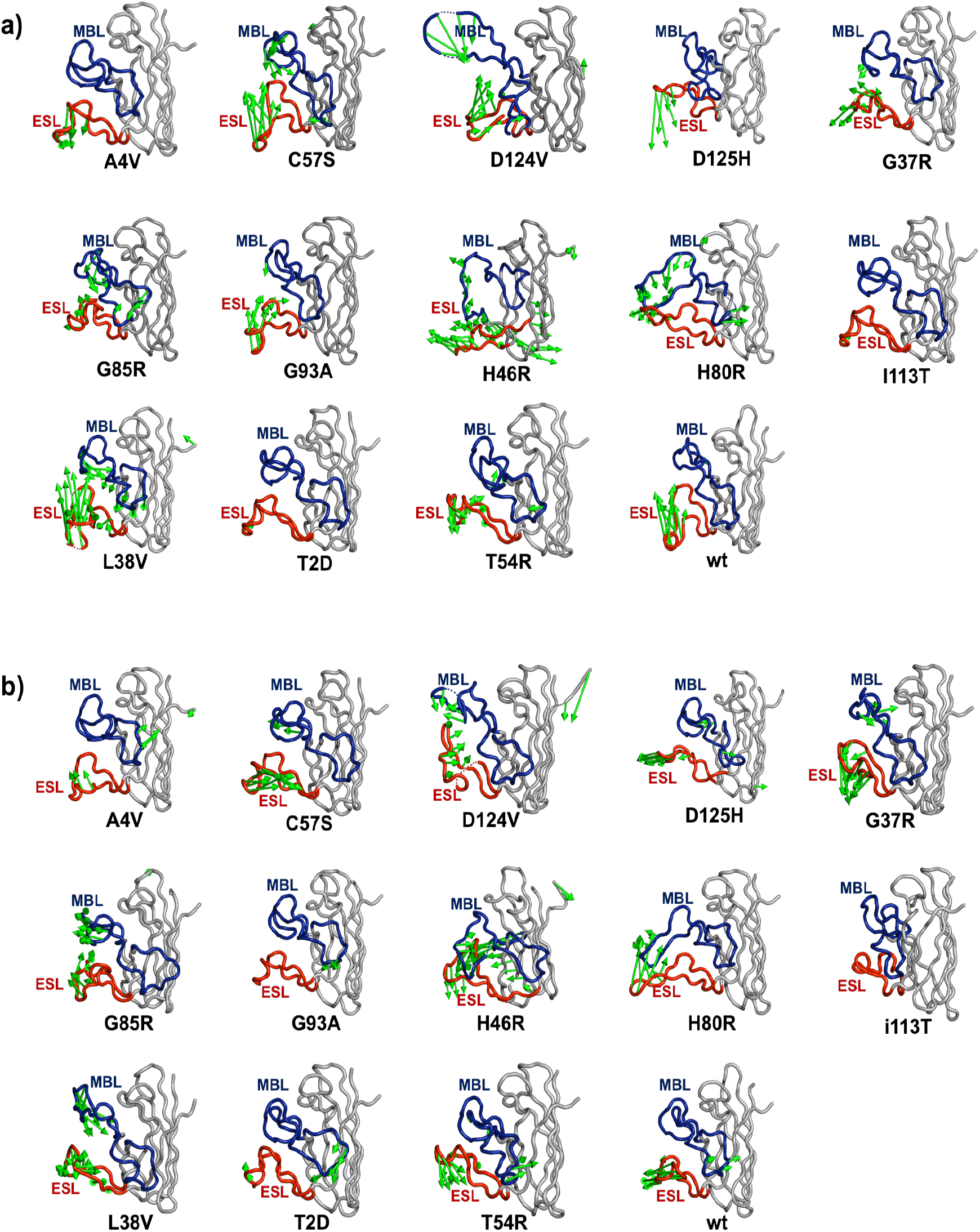
PCA analysis of the concatenated trajectories. Porcupine plots corresponding to the a) PC1 b) PC2. The wt-SOD1 and variant protein backbone is coloured in grey, the MBL in blue and ESL in red. The vectors coloured in green represent the direction and magnitude of the domain motion.

### Hydrogen bond analysis

To further investigate the dynamics of the wt-SOD1 and its variants we analysed the total number of hydrogen bonds formed during the simulations. The average number of hydrogen bonds formed in the entire SOD1 protein in the wt-SOD1, WTL and MBR variants are shown in Figure 7 (full distributions in Sup Fig 4). Interestingly, A4V and H46R variants, which commonly represented the two extreme opposites in the other analyses, had a similar number of hydrogen bonds as the wt-SOD1, while all other variants consistently presented a lower mean number with respect to the wt-SOD1.

**Figure 7.**
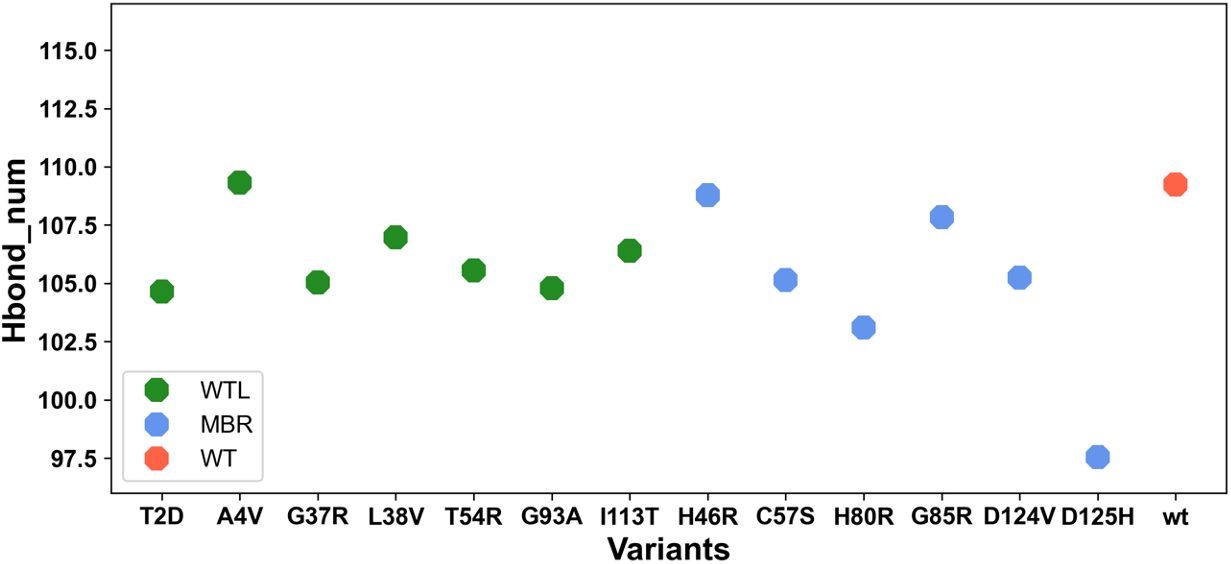
The mean number of hydrogen bonds formed in the wt-SOD1 (coral), WTL (green) and MBR (blue) variants during the entire length simulation.

### Graph-Theory based analysis

To better investigate the complex nature of the intramolecular interactions of the simulated systems, we used graph-theory based descriptors to evaluate the role of each residue within their interaction network. To this end, we used a weighted network representation, where each residue of the protein was a node of the network. Two nodes were connected by a link if the distance between their side chain centroids was lower than a threshold. In addition, each link was weighted with the contact frequency as calculated from the MD simulations trajectory. Several descriptors were computed using this network representation of the protein.

Many typical network descriptors (such as degree, strength, clustering coefficient, etc.) can capture local features of the system. Conversely, *shortest path based* descriptors are able to capture the connection between residues, even when they are spatially distant. Indeed, such quantities do not consider direct interactions but a set of consecutive interactions, defined as a path. For this purpose, we selected the closeness centrality descriptor, defined as the reciprocal of the sum of the length of the shortest paths between a given node and all other nodes in the graph. The higher the closeness of a residue, the higher its centrality is in the protein network.

First, we compared the distributions of residue closeness centrality values analysing wt-SOD1 and its 13 variant systems. We studied separately the residues belonging to metal binding loop and electrostatic loop. In Figure 8a, we reported the boxplots for each system, where the boxplots represent the values obtained considering all three simulation replicas.

**Figure 8.**
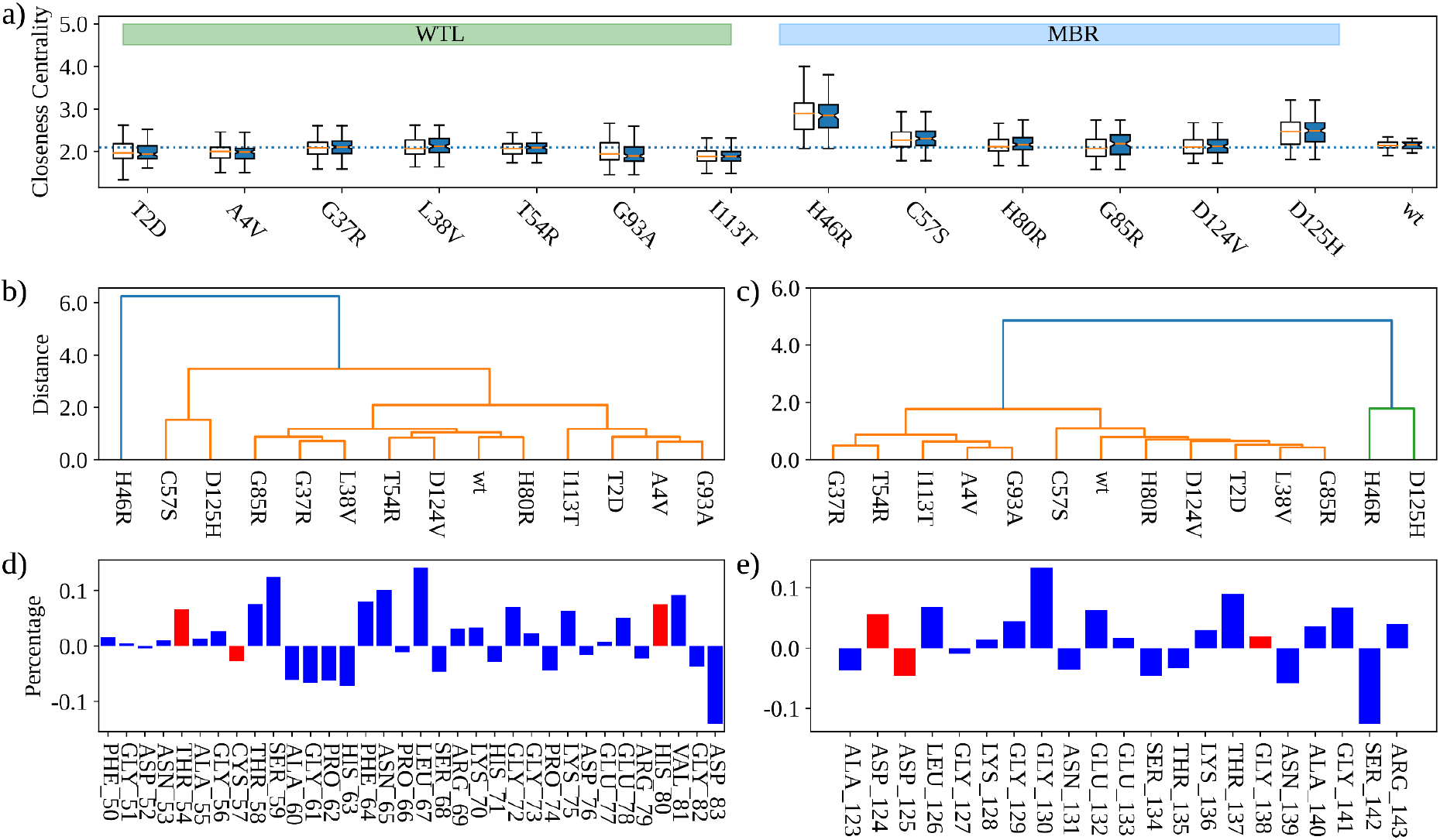
**a)** Box plots depicting Closeness centrality comparison of the wt-SOD1 and the variants. For each variant the boxplot on the left (respectively on the right) is obtained considering only the residues belonging to the MBL (resp. the ESL). The dotted line represents the mean Closeness centrality of the wt-SOD1 residues. **b)** Hierarchical clustering of the wt-SOD1 and its variants obtained considering the Closeness centrality indexes of the residues belonging to the MBL. **c)** Same as in c) but for the ESL residues. **d)** Mean difference (in percentage) between the closeness centrality indexes of the residues of the variants with respect to the wild-type ones for the MBL. **e)** Same as in d) but for the ESL.

Overall, the residue closeness centrality values of MBR variants were >= than the wt-SOD1 while WTL variants were <= than the wt-SOD1. H46R and D125H showed a large difference with respect to the wt system considering both metal binding loop and electrostatic loop regions (Figure 8a).

To further compare the wt-SOD1 with the variant systems, we performed a hierarchical clustering analysis (Figure 8b-c), which allowed us to compare each system with all the others. Figure 8b shows the clustering of the systems based on the centrality values of their metal binding loop residues. The metal binding loop region of H46R displayed the most singular behaviour, since it formed a group separated from all the others. When the variants were represented by the centrality values of the electrostatic loop residues, D125H and H46R formed a cluster distinct from all other variants (Figure 8c). In general, the clusters obtained using the metal binding loop and electrostatic loop residues (Figure 8b and Figure 8c) were very similar to each other, maintaining their global structure. In line with the previous results, H46R had a different profile from all the other systems while D125H clustered together with H46R based on metal binding loop residues and with C57S based on electrostatic loop residues.

Lastly, we investigated the contribution of each residue to the observed differences in the centrality values between variants and wt-SOD1. We calculated the mean percentage difference between the closeness centrality indexes of the residues, as calculated from the wt-SOD1 and the variants. The higher this value, the larger is the mismatch between the residues in the two classes of systems. We reported the results of this analysis in Figure 8d-e for metal binding loop and electrostatic loop regions respectively. The red bar represents residues that underwent a mutation in one of the simulated systems. Remarkably, the most marked differences did not regard mutated residues, testifying the complex nature of the interactions occurring in the protein.

### Covariance analysis

To study the difference between the dynamics of wt-SOD1 and its variants, we analysed the correlation between the covariance of the residue motion registered in such systems. For each MD simulation we built a covariance matrix calculating the covariance of the motion between each couple of residues. Hence, for each variant we computed the Pearson correlation coefficients between the covariance values regarding the wt and the three replicates of the variant systems. We reported in Figure 9a the mean and the standard error of the means of such correlations, where a high value means that the correlated motion between the wt and the variant systems was similar. T2D, I113R, C57S and G85R were characterised by a high mean correlation value, therefore highlighting an overall similar motion with the wt-SOD1. Conversely, once again H46R exhibits a marked difference with the wt-SOD1 behaviour.

**Figure 9.**
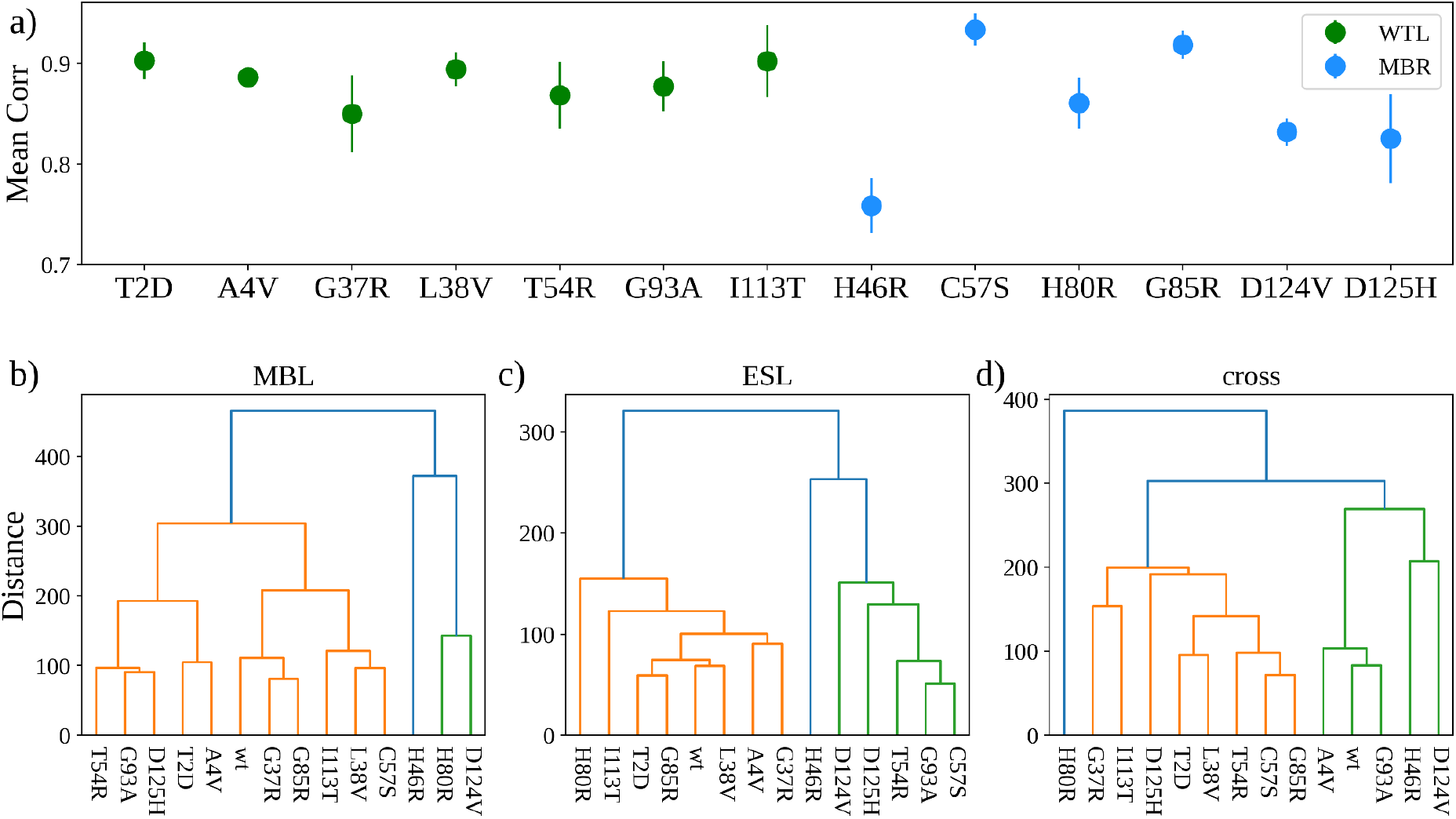
**a)** Mean correlation between the covariance matrices of the atomic coordinates of the wt-SOD1 and the variants. Bars represent the Standard Error of the Mean. Hierarchical clustering of the wt-SOD1 and ‘crystal’ variants obtained considering the covariances of the atomic coordinates of all the residues belonging to the **b)** MBL. **c)** ESL. **d**) covariances between the residues of the MBL and ESL.

In Figure 9b-d we showed the results of the hierarchical clustering using the covariance values as descriptors. Looking at panel b, c and d in Figure 9, variants such as G85R and G37R presented correlated motions similar to the wt-SOD1 when considering the two separately, while the cross covariances differed.

### Phenotype analysis of WTL and MBR variants

In order to investigate whether WTL and MBR variants associate with different clinical outcomes, we collected clinical data of patients carrying the variants investigated in this study. We were able to retrieve clinical data of 489 ALS patients with A4V, G37R, L38V, G93R, I113T, H46R, H80R, G85R and D125H (see details of individual variants in Table 4) [34]. Cox proportional hazard analysis (Figure 10) showed that patients with MBR variants presented a longer survival time (approximately 6 years median difference, p < 0.001) than patients with WTL variants, while no difference (p = 0.19) was observed for the age of onset (details in Table 4). Because A4V and I113T represented the great majority of the *SOD1* ALS patients in our sample (312 and 120 respectively), we repeated the survival analysis excluding these two variants from the WTL group. This analysis confirmed a significant difference between WTL variants (without A4V and I113T) and MBR variants (approx. 2.5 years median difference, p = 0.006).

**Table 4.** Characteristics of the clinical dataset.

|  | SOD1 variant | HGVS V2.0 nomenclature | Total sample size | Patients with disease duration | Number censored | Mean disease duration (months) | Median disease duration (months) | Patients with age of onset | Mean age of onset (years) | Median age of onset (years) |
| --- | --- | --- | --- | --- | --- | --- | --- | --- | --- | --- |
| 1. | A4V | A5V | 312 | 260 | 7 | 15.75 | 13 | 298 | 48.61 | 49.46 |
| 2. | D125H | D126H | 3 | 1 | 0 | 8.08 | 8.08 | 3 | 50 | 52 |
| 3. | G37R | G38R | 10 | 6 | 0 | 182.33 | 186 | 10 | 35.55 | 33.98 |
| 4. | G85R | G86R | 18 | 17 | 1 | 42.24 | 30 | 17 | 60 | 60 |
| 5. | G93R | G94R | 1 | 1 | 0 | 55 | 55 | 1 | 34 | 34 |
| 6. | H46R | H47R | 19 | 18 | 12 | 318.65 | 277 | 19 | 45.16 | 43 |
| 7. | H80R | H81R | 1 | 1 | 0 | 18 | 18 | 1 | 24 | 24 |
| 8. | I113T | I114T | 120 | 86 | 15 | 94.36 | 62 | 108 | 53.81 | 53 |
| 9. | L38R | L39R | 5 | 4 | 0 | 19.66 | 23.5 | 5 | 44.09 | 43 |
| 10. | All WTL | All WTL | 448 | 357 | 22 | 37.32 | 14 | 422 | 49.54 | 50 |
| 11. | All WTL (no A4V and I113T) | All WTL (no A5V and I114T) | 16 | 11 | 0 | 111.6 | 55 | 16 | 38.12 | 38.33 |
| 12. | All MBR | All MBR | 41 | 37 | 13 | 176.65 | 84.17 | 40 | 51.3 | 50 |

**Figure 10.**
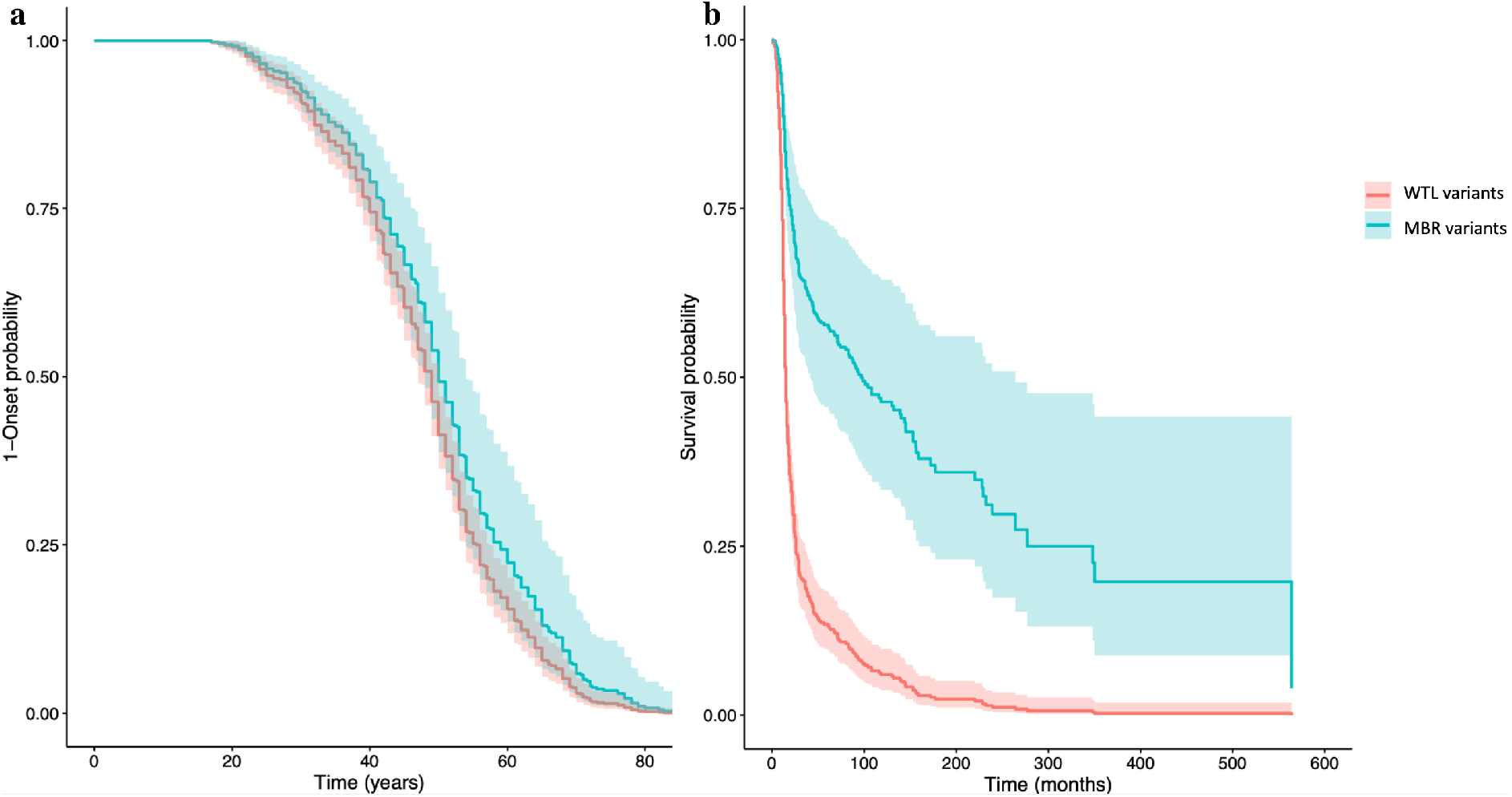
Age at onset (a) and survival (b) curves from the cox proportional hazard analysis (all WTL variants VS all MBR variants).

## DISCUSSION

In this study we investigated the differences in the structural and dynamic behaviours of the wt-SOD1 and sets of its WTL and MBR variants associated with ALS. Our results showed the disparate nature of dynamic properties of WTL and MBR variants. We showed that in the two groups there were differences in key features such as flexibility of the metal binding loop and electrostatic loop. It was shown previously that enervating electrostatic loop can lead to the gain of interaction, thereby increasing the formation of SOD1 amyloid fibrils [16, 38, 39]. Destabilization of the electrostatic loop due to disease causing mutations has long been linked to ALS pathogenesis [40, 41]. Akin to these previous studies, we also observed a substantial flexibility in the electrostatic loop in wt-SOD1, WTL and MBR variants. However, the electrostatic loop was substantially more flexible in the MBR variants in comparison to the WTL variants and wt-SOD1, which might be the reason why, in our simulations, MBR variants adopted more extended conformations with respect to wt-SOD1 and WTL variants as suggested by the radius of gyration analysis.

Our study also illustrated that H46R had the largest impact on flexibility, compactness of the protein structure, conformational changes, and domain motions. We also showed that there was a significant difference in the flexibility of the metal binding loop and electrostatic loop in A4V and H46R variants with respect to the wt-SOD1 and these two variants represented the two flexibility extremes across the variants that we studied. Interestingly, these two variants are known to represent extremes of the phenotypic spectrum of *SOD1* ALS. A4V variant causes the fast progression of ALS, with a survival time of 2 years [19–21] whilst H46R causes a slow progressing form of ALS, with a mean survival of ~12 years [22–25].

The disease-causing mutations in SOD1 protein can adopt a range of misfolded states [41] which are challenging to sample computationally. Our MD simulations showed that the MBR variants, C57S and G85R, adopted a range of conformational changes, while D124V, D125H and wt-SOD1 adopted one major conformation. In comparison, the WTL variants, T2D, G37R, adopted three conformational changes. The PCA performed on the trajectories of the simulations we performed showed that the range of conformational changes observed in the WTL and MBR variants were led by the domain motions in the electrostatic loop and metal binding loop regions. The motion and flexibility of the electrostatic loop were dynamic features observed ubiquitously in the SOD1 variants we studied.

The hydrogen bond analysis performed in our study, highlighted differences in the behaviour of the wt-SOD1, WTL and MBR at an atomic level. We observed that the largest number of hydrogen bonds were formed in the wt-SOD1 and A4V variant during the simulations. This is a very interesting observation, despite acting very similar to the wt-SOD1 on an atomic level the A4V variant manifests itself as a fast-progressing form of ALS. Previous studies have pointed out that the pathogenic mutations in SOD1 might alter the protein hydrogen bonds and promote the formation of new ones which can increase the probability of misfolding and aggregation in SOD1 [42, 43]. Our results complement these by showing that apart from A4V, SOD1 variants generally form fewer hydrogen bonds which might affect the protein stability and favour misfolding.

We also showed that the dynamic behaviour of the L38V variant was unexpected. Although prior studies have grouped L38V as a WTL variant [27] and L38V and wt-SOD1 have very similar crystal structures (RMSD = 0.074 Å), the difference between their dynamic behaviours was highly noticeable. In our study we observed that the increased flexibility and dominant motion of the electrostatic loop of L38V were similar to the MBR variants. This might be due to the disruption of the β-barrel plug [27]. Interestingly, L38V has been associated with early onset of ALS (mean age of onset = 38 years) [44].

Driven by the consistent differences between WTL and MBR variants highlighted in our analyses, and by the recurrent positioning of A4V and H46R at the far ends of such differences, mirroring the extreme clinical outcomes these two variants are associated with (fast and slow progressing forms of ALS), we attempted to investigate if WTL and MBR variants led to different clinical outcomes. Using a large clinical dataset (almost 500 *SOD1*-ALS patients) we were able to show that the survival of patients with a WTL variant was substantially shorter than the survival of patients with an MBR variant (median difference approximately 6 years, p < 0.001).

Experimental data suggests that ALS arises from a toxic gain-of-function mechanism and loss of SOD1 function alone has been linked to a severe phenotype distinct from ALS [45–47]. However, loss of function of SOD1 in ALS has been proposed as a potential modifier. A key difference between WTL and MBR variants is their effect on the SOD1 function. MBR variants, but not WTL variants, cause reduction of SOD1 enzymatic function in *in vitro* experiments of the isolated and engineered SOD1. On such a basis, it follows that our results support the hypothesis of decoupling between the role of gain and loss of SOD1 function in ALS affecting disease onset and duration respectively and potentially independently.

Identifying and understanding which mechanisms contribute to the development of the disease and which to its progression can improve genetic counselling, the development of new therapies and the design of more effective trials. This is particularly timely for *SOD1* given the current clinical trials that are testing the efficacy of treatments based on *SOD1* antisense oligonucleotides [48]. The great difference in the survival time between the patients carrying WTL and MBR variants, and the structural features that we found associated with the two classes, could be used to improve the classification into fast and slow progression of the patients that take part to trials, and to generate better estimates of their expected survival. A correct estimation of the expected survival is essential for the design of trials and for the interpretation of its results, as potential beneficial effects on slow-progressing variants would require lengthier trials to prove them.

A potential limitation of the design and interpretation of our study is the definition of the two classes of variants. The distinction between WTL and MBR variants was based on whether or not the variants were expected to affect the catalytic activity of SOD1. However, this effect was based on *in-vitro* studies of isolated and engineered proteins. An analysis of *SOD1* activity in blood samples from ALS patients showed that SOD1 activity was approximately halved in patients carrying a wide range of variants including some of the variants we classified as WTL [49]. Although these results are apparently in contrast with the *in vitro* studies we have based the classification of WTL and MBR variants on, a possible explanation could be that the loss of SOD1 function observed in the blood of patients is not an intrinsic effect of the variants but the result of a secondary effect linked to other disease pathogenic mechanisms, e.g. SOD1 aggregation. Another possibility is that both WTL and MBR variants affect SOD1 function but with different effect sizes.

In conclusion, our study highlights key differences in the dynamics of the WTL and MBR *SOD1* variants, and wt-SOD1. It sheds light into the behaviour of *SOD1* variants at an atomic and a molecular level, suggesting interesting structural features that could be investigated to explain their associated phenotypic variability, and it supports the role of loss of function of SOD1 as a modifier of disease progression in ALS.

## METHODS

### Protein structures

The protein structures of wt-SOD1 and 13 variants were obtained from Protein Data Bank (PDB) [50]; the detailed structural information is shown in Table 1. SOD1 variant structures were carefully selected; only those variants which had a single point mutation were included. An in-house python script Mutatpipe.py was utilized to select and the filter the variants (https://github.com/Utilon/MutaPipe_Repo). For the variants which had more than one crystal structure, the protein structure with the highest resolution and without missing residues was selected. SOD1 exists in a homodimer form in nature, however, for the scope of this study we focused only on the SOD1 monomer; the additional chains and the copper/zinc ions present in the PDB structures were manually removed. The missing residues in 1OEZ, 3H2Q, 3H2P and 1P1V monomer structures were modelled in with Modeller 10.2 program [51].

### Variants nomenclature

The old nomenclature was used to name *SOD1* variants as this reflects the actual position of the amino acids in the chains of the SOD1 structures we have used in our study. The new Human Genome Variation Society (HGVS) v2 nomenclature numbers the amino acids according to the mRNA reference sequence (GenBank: NM_000454.4) and it includes the start methionine which is cleaved post-translationally. Table 1 reports both nomenclatures for each variant.

### MD simulations protocol

We employed MD package GROMACS (version 2020.1) [52–54] to perform all atom MD simulations of wt-SOD1 protein, WTL and MBR variants. All the MD simulations were performed on the GPUs at *Rosalind* (https://rosalind.kcl.ac.uk), the high-performance computing (HPC) facility at King’s College London. AMBER99SB-ILDN forcefield was employed for all the MD simulations performed in the study [55]. The structure was solvated in the TIP3P water model in a rhombic dodecahedron box; the protein was placed at least 1.4 nm from the edges in all directions. Na^+^ and Cl^-^ ions were then added to neutralize the system and then the system was energy minimized over 50000 steps using the steepest descent method. The whole system consisted of ~ 30237 atoms; the number slightly varied in different variants.

After energy minimizing the system the solvent and ions around the protein were equilibrated in two steps. The first step was carried out under the NVT ensemble (constant number of particles, volume and temperature) for a length of 100 ps, at 300 K, using Berendsen thermostat. The second step was carried under NPT ensemble (constant number of particles, pressure and temperature), using the Parrinello-Rahman barostat [56] for a length of 100 ps at 1 atm and 300 K.

The final production was performed in triplicates for a length of 100 ns at 300 K for the 13 variants. In case of the wt-SOD1 the simulations were repeated 6 times for a length of 100 ns at 300 K. The covalent bonds were constrained using Linear constraint solver (LINCS) algorithm [57] and Particle Mesh Ewald (PME) algorithm was used to calculate long range interactions [58]. Leap frog integrator was used to calculate the equation of motion with a timestep of 2 fs [58]. The detailed information about the protein structures and total number of MD simulations performed in this study are summarized in Table 1.

### MD trajectory analysis

The visual analysis was performed using PyMOL [59] and Visual Molecular Dynamics (VMD) program [60]. We calculated the Cα RMSF, Cα RMSD, radius of gyration and hydrogen bonds using the GROMACS modules; *gmx rmsf, gmx rms, gmx gyrate, gmx hbond*.

### Principal Component Analysis

PCA is a highly used technique to reduce the dimensionality of the data obtained from molecular dynamics simulations and extract the dominant motion in the proteins [35, 36]. For PCA, the trajectories from three replicates were concatenated into one trajectory and the protein backbone was selected for analysis. *gmx covar* was then used to calculate and diagonalize the mass weighted covariance matrix. *gmx anaeig* was used to analyse the eigenvectors. The structures corresponding to PC1 and PC2 were extracted. To make the porcupine plots the structures corresponding to PC1 and PC2 were loaded to PyMOL. The vectors representing domain motion and direction were constructed after uploading the python script Modevectors.py on PyMOL (https://pymolwiki.org/index.php/Modevectors).

### Statistical Analysis

All the statistical analyses in the study were performed using SciPy.Stats computing library in Python [61]. Wilcoxon Rank-sum *T*est was employed to calculate the differences in the statistical significance of the RMSF between the variants and the wt-SOD1. Age of onset and survival time from disease onset analyses were performed using Cox proportional hazards regression. Models were adjusted for site of onset of symptoms and sex. The coxph() function was utilised with tie resolution at the default setting. When modelling survival time from onset, age of onset was also included as a covariate. For all statistical analyses, p < 0.05 was considered significant.

### Graph theory analysis

We modelled each protein structure as a network, schematizing each residue as a node of the network and each intramolecular interaction as a network edge [62, 63]. In particular, we defined a weighted graph for each system, weighing each link connecting two residues with the corresponding contact frequency calculated from the molecular dynamics, defining, in this case, contact between two residues if their distance is less than 8.5 A, similar to previously described procedures [64].

### Covariance Analysis

Covariance matrices of the atomic positions of all the atoms of the metal binding loop and electrostatic loop were computed from molecular dynamics trajectories. Covariance between couples of residues were then obtained averaging the covariances of all the couples of atoms belonging to the considered residue couple.

### Clinical dataset

Data on people with *SOD1*-ALS were collected from the ALS Online Database (https://alsod.ac.uk) [7, 8], the Project MinE whole-genome sequencing dataset [65], and a number of centres. The corresponding authors of the AlSoD database entries with missing data were contacted to fill the gaps. Anonymised records of people with *SOD1*-ALS were obtained from the following centres: Macquarie University, ANZAC Research Institute, University of Massachusetts, University Hospitals of Montpellier, King’s College London, Washington University School of Medicine in St Louis, Peking University Third Hospital, Università degli Studi di Siena, Northwestern Medicine – Feinberg School of Medicine, Istituto Auxologico Italiano IRCCS-University of Milan, University of Belgrade and University of Calabria. Individuals were eligible to be included if their diagnosis of ALS was made by a consultant neurologist, or their data was reported in the literature with an ALS diagnosis. The clinical and demographic features utilised in this study were country of origin, sex at birth, age of onset (in years) of first motor symptoms, site of onset, survival time (in months) defined as time from diagnosis to death or latest visit. In each analysis, individuals with missing data among the subset of clinical features required were discarded. Individuals with the same *SOD1* variant, country of origin, sex, age and site of onset were considered to be duplicates.

## ACKNOWLEDGEMENTS

We acknowledge use of the research computing facility at King’s College London, Rosalind (https://rosalind.kcl.ac.uk), which is delivered in partnership with the National Institute for Health Research (NIHR) Biomedical Research Centres at South London & Maudsley and Guy’s & St. Thomas’ NHS Foundation Trusts and part-funded by capital equipment grants from the Maudsley Charity (award 980) and Guy’s and St Thomas’ Charity (TR130505). The views expressed are those of the author(s) and not necessarily those of the NHS, the NIHR, King’s College London, or the Department of Health and Social Care. We would like to thank people with ALS and their families for their participation in this project. We thank the Project MinE Sequencing Consortium members for their collaboration and support.

## APPENDIX 1 Project MinE ALS Sequencing Consortium group authors

Philip van Damme, Philippe Corcia, Philippe Couratier, Orla Hardiman, Russell McLaughlin, Marc Gotkine, Vivian Drory, Vincenzo Silani, Nicola Ticozzi, Jan H Veldink, Leonard H van den Berg, Mamede Carvalho, Susana Pinto, Jesus Mora Pardina, Monica Povedano, Peter M Andersen, Markus Weber, Nazli Başak, Ammar Al-Chalabi, Chris Shaw, Pamela Shaw, Jonathan Cooper-Knock, Alfredo Iacoangeli, Karen Morrison, John Landers, Jonathan Glass, Patrick Vourc’h.

