## Supplementary figures for "Molecular dynamics analysis of Superoxide Dismutase 1 mutations suggests decoupling between mechanisms underlying ALS onset and progression"

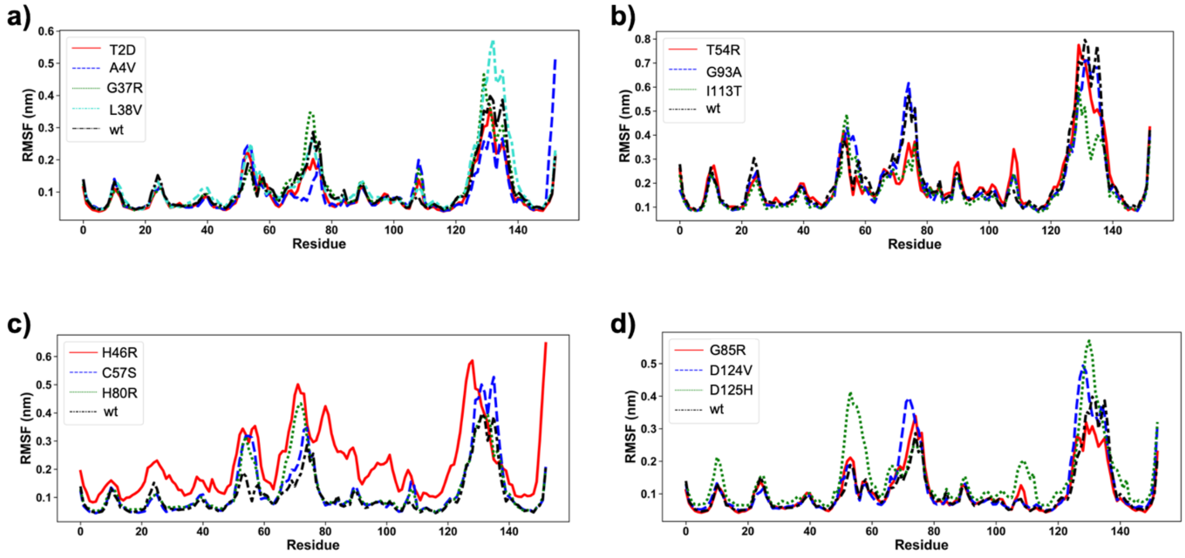


**Sup. Figure 1** RMSF comparison of the entire length of the SOD1 protein in case of a) & b) WTL variants and c) & d) MBL variants.

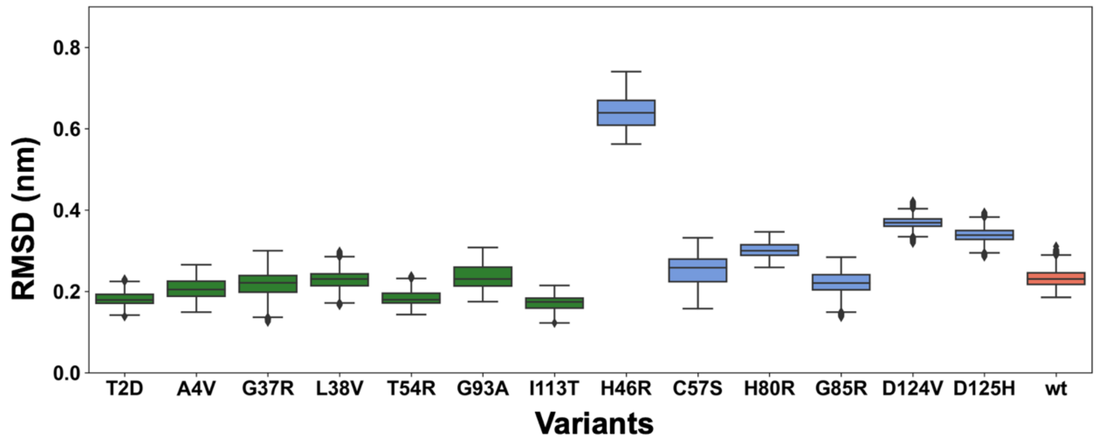


**Sup. Figure 2** Box plots depicting RMSD analysis of the MD simulations after reaching the equilibration phase (>20ns) performed on the wt-SOD1 (coral), WTL (green) and MBR (blue) variants. The RMSD is calculated for the Cα atoms of the wt-SOD1 and the variants. The whisker bars represent the range of minimum and maximum RMSD, the median is represented by a line subdividing the box.

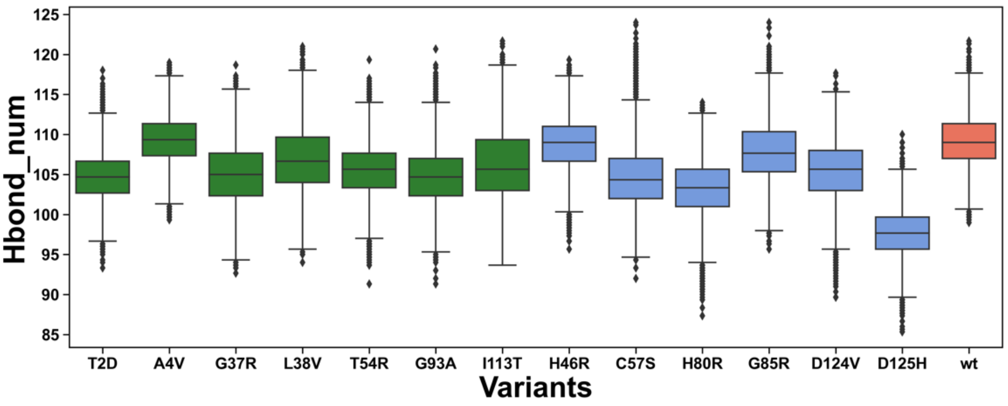


**Sup. Figure 3** Box plots depicting of the Hydrogen bond distribution of the wt-SOD1 (coral), WTL (green) and MBR (blue) variants.
